# Molecular dynamics of membrane partitioning of nifedipine prior to binding calcium channel Cav1.1

**DOI:** 10.1101/2025.03.27.645691

**Authors:** Warre Dhondt, Louis Vanduyfhuys, Ahmad Reza Mehdipour

## Abstract

An increasing number of ligand-bound membrane protein structures reveal ligand-binding sites on the lipid-exposed surface of the protein within the membrane bilayer. Binding events to such sites have previously been studied using molecular dynamics (MD) simulations and experiments in cases such as calcium-gated potassium channels^1^ and sodium channels^2^. The proposed binding mechanism is that these ligands partition into the membrane to gain access to their binding site. What is currently unavailable is what the thermodynamic and kinetic contributions of the ligand-membrane and ligand-protein interactions are to the overall binding event. Here, we used MD simulations and enhanced sampling methods to study the membrane partitioning of a DHP calcium channel antagonist, nifedipine, from the voltage-gated calcium channel Cav1.1. We present that drug-membrane interactions occur on a much faster timescale than the overall binding of nifedipine to Cav1.1.

## Introduction

Voltage-gated ion channels play a key role in the pathophysiology of cardiovascular disease and pain, which makes them an attractive target for therapeutic intervention. Ion channel modulators are traditionally divided into two categories: pore blockers, which obstruct the flow of ions through the central pore, and allosteric modulators, which bind outside the pore to stabilize a specific conformation in the functional cycle of ion channels. The link between the mechanism of action of these classes of inhibitors and their selectivity was first established for the block of sodium channels by local anesthetics^3–5^. It was found that the inhibition of sodium currents depends on both the rate of membrane depolarization and the holding membrane potential, with different dependencies for different classes of inhibitors. The blockade by a given inhibitor would either be enhanced by a greater depolarization frequency, or by lower membrane potentials, or both. The resulting “modulated receptor hypothesis” invokes alternative drug access routes to explain this phenomenon^3^: polar compounds would enter the open pore from the solvent, whereas hydrophobic compounds would bind from within the membrane compartment. This observation raises the question of how the latter class of inhibitors reach their binding site from inside the membrane. The proposed “membrane access pathway” rationalizes how binding within the membrane compartment may occur^6,7^. For hydrophobic drugs, partitioning into an equally hydrophobic environment, such as the hydrocarbon core of a membrane bilayer, would be energetically favorable. From this location, the drug could then access the binding site on the transmembrane region of the target protein.

Drug binding and dissociation pathways leading into the membrane have been experimentally determined for several systems. Mutagenesis studies where the channel opening into the membrane was expanded or constricted, combined with inhibition experiments, showed that drug entry into the central pore via inter-subunit fenestrations is essential for blocking closed Nav, BK, and NDMAR ion channels^1,8,9^. Such binding paths have also been described for the class A GPCRs such as Rhodopsin^10^, the β_2_-adrenergic receptor^11^, and the melatonin receptor^12^. Molecular dynamics (MD) simulations have been used to chart the binding pathway of pore blockers from the membrane to the pore of closed ion channels to atomic detail^13–15^. However, these studies focused on how the drug may pass through the inter-subunit fenestrations rather than the biophysical nature of this class of binding event. This leaves several factors that appear to be highly relevant to the thermodynamics and kinetics of binding, including the role of the membrane in this binding mechanism. By considering this binding pathway as a two-step binding mechanism consisting of the partitioning into the membrane and subsequent binding to the receptor, it can be seen that the drug-membrane interactions will contribute to the overall binding affinity and binding kinetics^16^. Decomposing the overall binding event into the contribution of drug partitioning and drug binding, both in terms of thermodynamics and kinetics, is interesting for understanding the mechanism of action for this class of compounds. In the present work, we have used MD simulations in combination with enhanced sampling to study how nifedipine, a prototypical dihydropyridine (DHP) antagonist of Cav1.1, interacts with a POPC membrane and how this drug-membrane interaction compares to the overall binding thermodynamics and kinetics.

## Results

### Unbiased MD simulations of nifedipine-membrane interactions

For nifedipine to reach its membrane-buried binding site on Cav1.1, it must first interact spontaneously with the membrane lipids. Therefore, we set up unbiased MD simulations to study the interaction between nifedipine and a model phospholipid membrane comprising 100% POPC lipids. The membrane model was validated by comparing the structure, packing, and dynamics of the lipids to experimental measurements to good agreement (Table 1).

**Table 1:** Calculated versus experimental parameters of POPC membranes. The area per lipid (APL), bilayer thickness (D), and lateral diffusion coefficients (D_xy_) were calculated based on an MD simulation of a 7 nm x 7 nm POPC membrane presented in this study (indicated in bold) compared to experimental measurements^17–21^. Temperature is indicated between brackets.

| APL ($\text{\AA}^2$) | D ($\text{\AA}$) | $D_{xy}$ ($10^{-7} \text{ cm}^2\text{s}^{-1}$) |
| --- | --- | --- |
| **$64.6 \pm 0.1$** | **$38.8 \pm 1.5$** | **$1.465 \pm 0.23$** |
| $68.3 \pm 1.5$ (303 K)<sup>16</sup> | $38.2$ (303 K)<sup>16</sup> | $0.8\text{-}1.6$ (296 K)<sup>17</sup> |
| $64.3 \pm 1.3$ (303 K)<sup>17</sup> | $39.1 \pm 0.78$ (303 K)<sup>17</sup> | $1.9 \pm 0.1$ (322 K)<sup>18</sup> |
| | | $1\text{-}2$ (303-323 K)<sup>19</sup> |
| | | $1.2\text{-}1.5$ (308, 313 K)<sup>20</sup> |

In four simulations where nifedipine was placed in the water layer above the membrane, the ligand spontaneously interacted with the lipids and entered the membrane for a prolonged time (>50 ns) in all the replicates within 800 ns (Figure 1a-b). During this interaction, nifedipine’s average position was ∼14 Å from the bilayer center in the transition zone between the polar and hydrophobic regions of the membrane (Figure 1b-c). Here, the amphipathic structure of nifedipine would be optimally satisfied by allowing the polar DHP ring to interact with the polar headgroups while positioning the nitrophenyl ring towards the hydrophobic core (Figure 1c). Next, we investigated whether the interaction with the membrane lipids affects the conformation of the ligand. The only appreciable internal coordinate of nifedipine is the dihedral angle between its two rings, χ_c8, c11-13_ (Figure 1d, inset). In solvent, the dominant conformation is one where the nitro group of nifedipine can interact and aligns with the methyl esters on the DHP ring. This state remains populated after partitioning into the membrane, but an additional intermediate state at χ_c8, c11-13_=50-60° becomes stabilized where the nitro group is between the methyl esters (Figure 1d). Because this angle resembles the bound pose of nifedipine bound to Cav1.1, the same dihedral angle was compared to simulations of nifedipine bound to Cav1.1^22^. Indeed, the rotation of this dihedral angle is restricted in the bound pose, and is similar to the state stabilized while interacting with the lipids (χ_c8, c11-13_=50-60°, Figure 1d). The interactions between nifedipine and the membrane may therefore stabilize the receptor-bound conformation.

**Figure 1:**
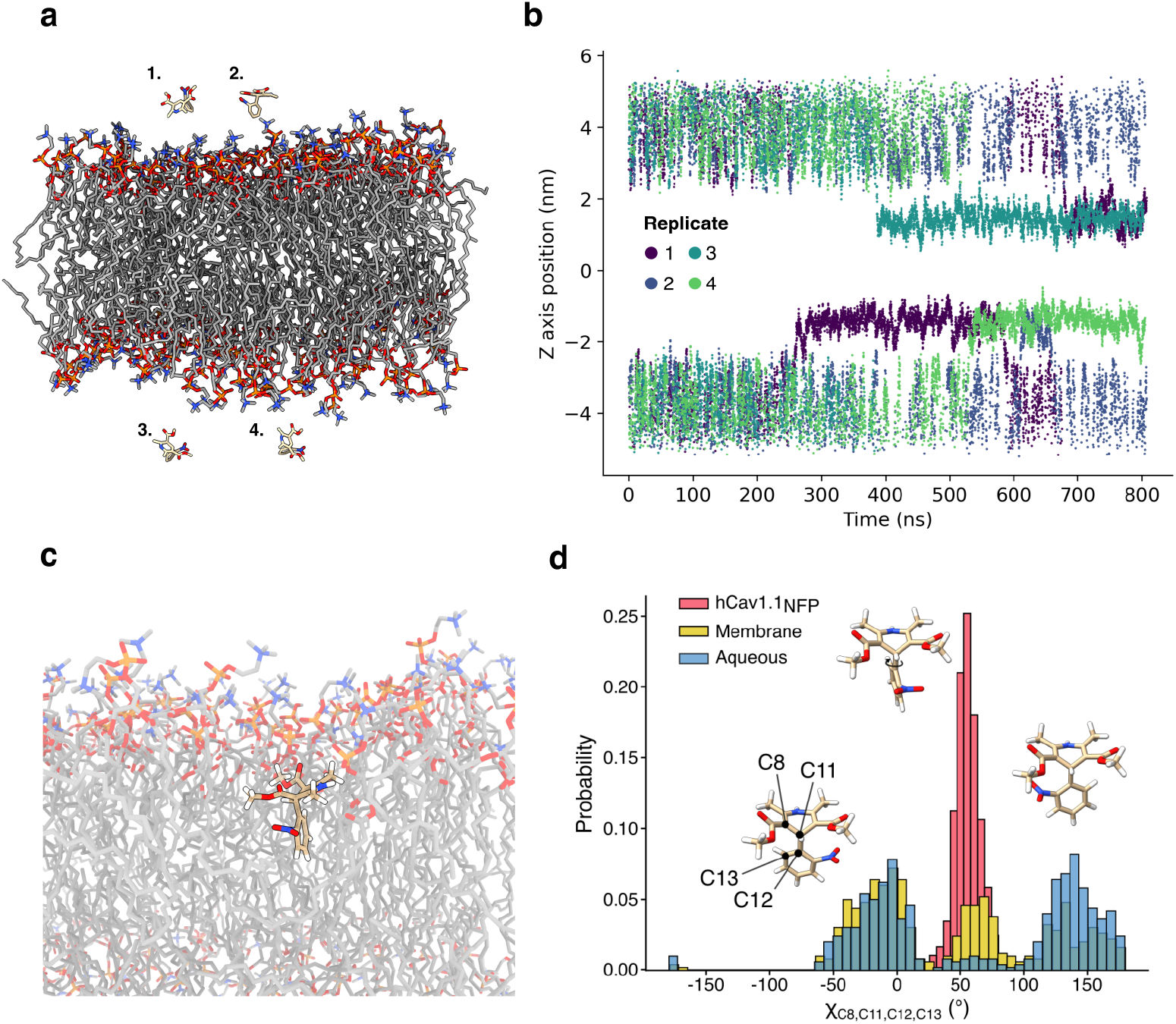
Spontaneous interactions between nifedipine and a model membrane. a) A representative model for the simulation setup that was used to study spontaneous interactions between nifedipine and a POPC membrane. b) Distance of the center of mass of nifedipine from the center of the membrane during four replicate simulations of the setup shown in a). c) A representative snapshot of nifedipine after membrane partitioning is located near the lipid headgroups. d) Probability densities for the dihedral angle χ_c8, c11-13_ of nifedipine across simulation frames where the ligand is in the water layer, inside the membrane, and when the ligand is bound to Cav1.1.

### Enhanced sampling of nifedipine-membrane interactions

To quantitatively describe the permeation of nifedipine through a model membrane, umbrella sampling simulations were performed along the permeation coordinate *d*_*z*_ (Figure 2a). Because the unbiased simulations only provided initial configurations for distances up to *d*_*z*_ =7 Å, additional simulations were performed in which the ligand was gradually pulled across the membrane with a moving potential at the ligand’s center of mass. The snapshots were then used as a starting point for umbrella sampling. The resulting free energy profile (FEP) was unexpectedly asymmetric with respect to the center of the membrane (Figure 2b-c). The FEP for this class of system is expected to be centrosymmetric due to the symmetry of the bilayer, whereas this profile reveals that the lower leaflet would be disfavored by approximately 40 kJ/mol. This phenomenon persisted regardless of the reconstruction algorithm applied to combine the different umbrella windows or the windows’ overlap (Figure 2b-c). In agreement with other work^23^, we found that the non-equilibrium pulling simulations introduced large deformations in the membrane that did not relax on the simulated timescales, which resulted in an artificial preference for leaflet whose headgroups the ligand was interacting with, and thus the asymmetric FEP (Figure 2d). This was confirmed by the fact that reversing the pulling direction during the simulation reverses the membrane asymmetry (Figure 2e).

**Figure 2:**
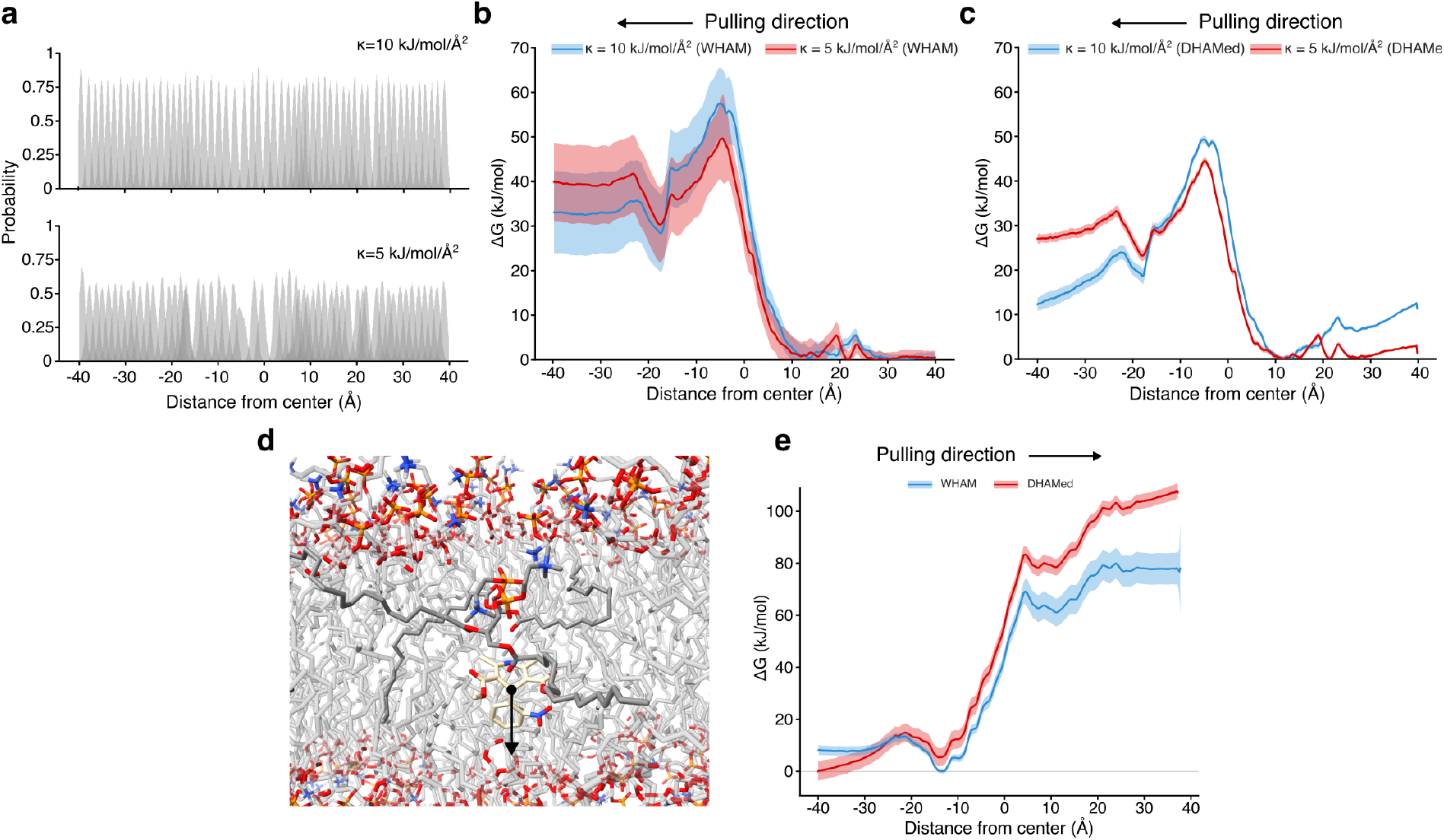
Introduction of structural artifacts by steered MD simulations. Nifedipine was pulled across a POPC membrane to generate starting snapshots for umbrella sampling. a) Histograms showing the sampling of the position of nifedipine along *d*_*z*_ over 57 umbrellas using different harmonic force constants. b, c, e) free energy profiles of nifedipine permeation through the POPC bilayer reconstructed based on umbrella sampling using steered MD frames as starting configurations. The FEPs were reconstructed using WHAM or DHAMed, and the pulling direction is indicated. d) snapshot for *d*_*z*_=-12 Å showcasing a POPC lipid being dragged along with the ligand during the steered MD simulation.

As the next step, we used shooting moves to sample transitions across the hydrophobic membrane core (see Methods). The resulting FEP shows local minima near the headgroup/hydrocarbon transition zone, supporting the observed spontaneous drug-membrane interactions, as well as a global maximum near *d*_*z*_ =0 where the ligand must cross the hydrocarbon core to move between the opposing leaflets (Figure 3b). In line with prior work^24,25^, we observe that the transition across the hydrocarbon core involved a flipping motion, quantified by the tilt angle θ (Figure 3a). When the ligand crosses the center of the membrane, we observe a 180° rotation in the preferred tilt angle such that the ligand aligns its amphiphilic axis with that of the lipids, as observed in the unbiased simulations (Figure 1c). This process is analogous to the so-called “flip-flop” of membrane lipids.

**Figure 3:**
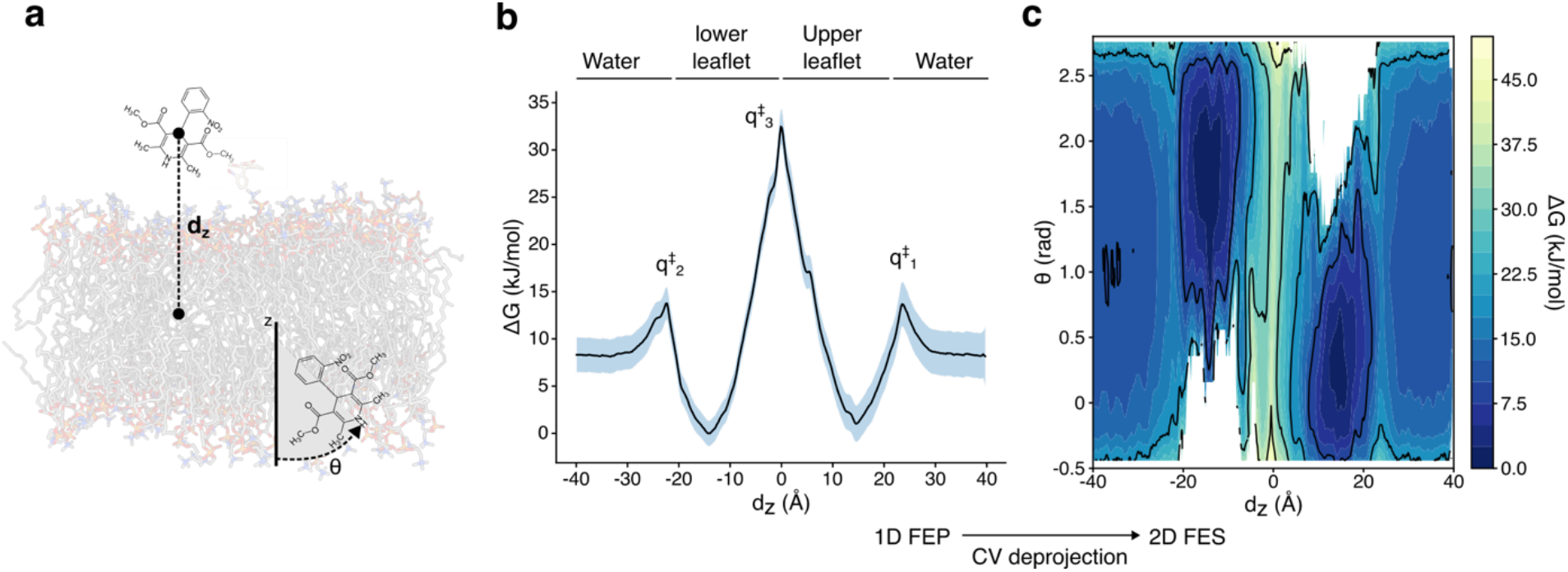
Free energy profile describing the permeation of nifedipine through a POPC bilayer. a) Description of the two collective variables (CVs) used to describe the permeation process. The first CV, d_z_, describes the distance between the ligand center of mass and that of the bilayer. The second CV, theta, describes the tilt between the amphiphilic axis of the ligand and the membrane normal. b) Free energy profile of nifedipine permeation after performing umbrella sampling along d_z_. The water layers and membrane leaflets, as well as local maxima (q^‡^) are indicated. c) The 1D FEP along d_z_ was transformed to a 2D free energy surface (FES) along d_z_ and theta.

A quantitative interpretation of this FEP requires considering that free energy differences between the aqueous state and the membrane-interacting state are dependent on the size of the water layer during the simulations. Increasing the volume of the water layer entropically stabilizes the aqueous state while leaving the membrane-interacting state unaltered. This effect is different from a change in drug concentration due to altering the x and y dimensions of the simulation box. In that case, the volume of the membrane phase will increase proportionally, maintaining the same partitioning coefficient/equilibrium constant (K). Here, the simulations were performed at a ligand concentration of ∼10 µM and a water layer of 25 Å. The mean energy difference between the membrane-interacting and aqueous states was similar for both leaflets at approximately ΔG=-5-6 kJ/mol in favor of the membrane-interacting state (Table 2). Calculated rates for membrane partitioning indicate that this event is ∼4 orders of magnitude faster than membrane permeation (Table 2). The experimentally determined binding rate of nifedipine to Cav1.1^26^ adjusted to the same concentration is 44.7 s^-1^ (experimental k_on_=4.47 x10^6^ M^-1^s^-1^), which is 4 to 5 orders of magnitude slower than the transition rates between leaflets (Table 2). Membrane partitioning and permeation would occur on a much faster timescale than the overall binding event, such that the ligand could reach its target from either local minimum inside the membrane.

**Table 2:** Free energy differences and rate constants for the permeation of nifedipine through a POPC bilayer. ΔG, free energy differences between the indicated macrostates are defined as the states between local maxima (see Figure 2). k, calculated rate constant for the transition between the indicated macrostates. 95% confidence intervals (mean ± 2 S.E.M.) are given between brackets.

| $\Delta G$ (kJ/mol) | $\Delta G$ (kJ/mol) | $k$ ($10^8$ s$^{-1}$) | $k$ ($10^8$ s$^{-1}$) | $k$ ($10^8$ s$^{-1}$) |
| --- | --- | --- | --- | --- |
| water→lower leaflet | water→upper leaflet | water→lower leaflet | water→upper leaflet | lower→upper leaflet |
| -5.802 | -5.161 | 57.047 | 70.256 | 0.006 |
| (-8.035; -3.569) | (-7.862; -2.460) | (17.605; 141.03) | (16.849; 199.86) | (0.002; 0.013) |
| | | $k$ ($10^8$ s$^{-1}$) | $k$ ($10^8$ s$^{-1}$) | $k$ ($10^8$ s$^{-1}$) |
| | | lower leaflet→water | upper leaflet→water | upper →lower leaflet |
|  |  | 5.706 | 9.269 | 0.006 |
|  |  | (2.240; 12.157) | (2.313; 25.818) | (0.002; 0.023) |

## Discussion

The unbiased MD simulations and free energy profile in this study show that nifedipine prefers to be located close to the lipid headgroups. The local minimum near z=±15 agrees well with the preferred locations of the related DHPs Bay K-8644 and amlodipine determined experimentally via x-ray scattering^27,28^. Although a first comparison with experimentally determined partition coefficients of nifedipine suggests that the calculated partitioning free energy is 15 kJ/mol lower than expected^6,29^, a quantitative comparison is not possible due to the geometry difference between the simulation box and the experimental setup. Comparison of the calculated rates of membrane partitioning and interleaflet flip-flop of nifedipine with the overall binding rates of nifedipine to Cav1.1 suggests that the rate-limiting step of binding is the interaction between nifedipine and the target protein, whereas drug-membrane interactions occur orders of magnitude faster. To further quantify the contribution of the membrane to the observed affinity, a study comparing nifedipine-membrane and nifedipine-protein interactions at biologically relevant concentrations is required.

## Methods

The equilibration and simulation conditions (except for duration of the simulation) were the same for all simulations. All simulations described next were performed using the MD engine GROMACS using the Charmm36m force field^30^. The ligand was parameterized using the Charmm general force field^31^. The systems were energy optimized for 5000 steps using steepest descent and subsequently equilibrated for 1 ns in NVT and 8.5 ns in an isothermal-isobaric ensemble at 310 K under periodic boundary conditions. Initial position restraints on the protein backbone and sidechain atoms and/or lipid headgroup atoms, as well as dihedral angle restraints were gradually released over the course of the equilibration to 0 at the start of the production run. Temperature and pressure were controlled during equilibration using the Berendsen thermostat and semi-isotropic barostat. Long-range electrostatic interactions were treated using Particle-Mash Ewald summation using 0.12 nm spacing and quartic interpolation. Integration was performed with a 2 fs timestep, and all H-X bonds were treated as rigid. The Berendsen thermostat and barostat were switched to Nose-Hoover thermostat and Parinello-Rahman barostat for the production runs to sample from the NPT ensemble.

### Modeling and unbiased MD simulations of nifedipine-membrane interactions

A unit cell containing a 75 Å x 75 Å patch of 1-palmitoyl-2-oleoyl-sn-glycero-3-phosphocholine (POPC) lipids (166 lipids) solvated by TIP3 water and 150 mM NaCl was constructed in CHARMM-GUI^32^ and simulated for 200 ns. To validate the membrane model, the area per lipid (APL), membrane thickness (D) and lipid diffusion coefficients were calculated (D_xy_). The mean APL was calculated by dividing the number of lipids by the instantaneous box size; the mean membrane thickness was calculated by averaging the distance between phosphorus atoms in opposing leaflets in each frame, then averaging over all frames. Uncorrected diffusion coefficients were calculated for two box sizes (75 Å x 75 Å and 50 Å x 50 Å) using the Python package DiffusionGLS after using the NPT unwrapping scheme proposed by von Bülow et al.^33^ implemented in MDAnalysis. The finite size effects were then calculated by using the Os^22^een correction implemented in *memdiff*^34^. Four frames were then extracted from the last 50 ns of the production run and nifedipine was placed in the solvent above the membrane patch, approx. 1 nm above the lipid headgroups (z = 3.5, z = 8.5) with different x and y positions and random orientations using *gmx insert-molecules*. The same force field parametrization for nifedipine as described above was used. The resulting systems were again energy optimized and equilibrated with initial position restraints, followed by 800 ns of unrestrained production MD in the NPT ensemble. To analyse the membrane partitioning of nifedipine, the distance between the center of mass (COM) of nifedipine (after unwrapping) and the COM of POPC residues was calculated.

### Modeling and MD simulations of hCav1.1

The cryo-EM model of the rabbit Cav1.1 bound to nifedipine was used to construct a homology model of the pore domain of human Cav1.1 (Uniprot ID Q13698). Based on rabbit Cav1.1α (PDB ID 6JP5), 20 candidate models for hCav1.1 were constructed using Modeller^35^. Candidate models of nifedipine-bound hCav1.1 with the highest DOPE quality score were used as a starting point for further editing. The loops between VSDI-VSDII (residues 377-417) and VSDII-VSDIII (688-787) are unstructured and were removed from the model. The protonation state of titratable side chains at pH 7.4 were determined using PROPKA 3^36^, except for the negatively charged residues of the ion selectivity filter (Glu292, 614, 1014, 1323), which were set as deprotonated. Disulfide bonds between Cys226-265, Cys245-261, Cys957-968 and Cys1338-1352 were applied. Two coordinated calcium ions from rabbit Cav1.1 were rigid body copied into the homology model as they are known to be essential for DHP binding^37^. The resulting structures were embedded in a homogenous (POPC) bilayer (160Å x 160Å), solvated by TIP3 water and 150mm NaCl in CHARMM-GUI^32^. The system was then equilibrated as outlined above and simulated for 500 ns.

### Steered molecular dynamics of nifedipine permeation

Starting from the final frames of the unbiased partitioning simulations, steered molecular dynamics (SMD) was used to pull the ligand across the remainder of the membrane. A harmonic potential of 50kJ/mol/Å^2^ centered on the COM of the ligand moving at a velocity of 0.1Å/ns was used to gradually move the ligand across the membrane from z=8 to z=40. These frames were then combined with those from the spontaneous partitioning event as starting configurations for umbrella sampling. Umbrella sampling was performed using the cartesian z-component of the distance between the COM of nifedipine and the COM of the POPC lipids as CV. The 57 starting configurations were spaced approximately 1.5 Å apart and sampled using 5 or 10 kJ/mol/Å^2^ harmonic force constants for 10 ns per umbrella. The different umbrellas were combined using a maximum-likelihood implementation of the weighted histogram analysis method (WHAM) in an in-house python package ThermoLIB. The FEP was also constructed using DHAMed implemented in the python package pyDHAMed (github.com/bio-phys/PyDHAMed). To estimate the errors for DHAMed, 10 independent reconstructions on bootstrapped samples were performed.

### Sampling nifedipine crossing of the membrane

To generate starting frames for umbrella sampling of nifedipine permeation across the membrane core, a ligand molecule was manually inserted at z=0 in a well equilibrated 75 Å x 75 Å patch of POPC lipids (see above). The resulting system was energy minimized and equilibrated with a 10 kJ/mol/Å^2^ harmonic potential on the ligand position. This was followed by 20 replicates of 5 ns of unrestrained simulation with randomized initial velocities.

### Umbrella sampling simulations of nifedipine-membrane permeation

Frames from the membrane-crossing transition paths and the unbiased simulations of nifedipine-membrane interactions were combined and used as starting configurations for umbrella sampling. In total, 80 windows spaced approx. 1 Å apart were simulated for 10 ns, writing the z-position to file every 0.2 ps. The different umbrellas were combined using a maximum-likelihood implementation of the weighted histogram analysis methods (WHAM) in an in-house python package ThermoLIB. Standard errors for the FEP obtained via WHAM were extracted from the variance-covariance matrix, which is calculated as the inverse of the Fisher information matrix of the MLE. Autocorrelation of the biased variable was accounted for by correcting for the statistical inefficiency in the variance estimate.

### Deprojection of free energy profiles

To gain more insight into the reaction pathways, 1D FEPs of membrane permeation were deprojected in terms of the permeation coordinate z, towards a 2D FES in terms of an additional collective variable. For this CV, the angle (theta) between the vector formed by C11 and C15 of nifedipine and the z axis, calculated using the Plumed CV ZANGLES. Such deprojection from a single CV *q* to 2 CVs (*q*_1_, *q*_2_) is based on Bayes’ theorem and can be expressed as follows^38^:

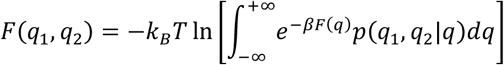

Herein, *p*(*q*_1_, *q*_2_|*q*) plays the key role and encodes the conditional probability to find the system in a state of given (*q*_1_, *q*_2_) on the condition that it already was in a state of given *q*. In the current investigation, the context is slightly different as we want to keep the original CV as one of the new CVs, i.e. we choose *q*_1_ = *q*, in this case we find:

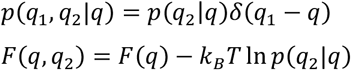

With the choice *q* = *z* and *q*_2_ = *θ*, we finally arrive at:

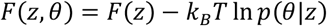

Herein, *p*(*θ*|*z*) now encodes the conditional probability to find the system at an angle of *θ*, given it already was at position *z*, which can be extracted from the performed molecular dynamics simulations. *F*(*z*) is the original 1D free energy profile constructed through umbrella sampling.

### Calculation of free energy differences and rate constants

The analysis of the free energy profiles to calculate free energy differences proceeded as follows. The macrostates were defined as all microstates between two local maxima. The free energy of a macrostate was calculated by integrating the probability distribution over all microstates contributing to that macrostate, then taking the negative logarithm and multiplying by k_B_T. Free energy differences were calculated by subtracting the free energies of macrostates. Rate constants were calculated based on transition state theory where the pre-factor was approximated as in Dubbeldam et al^39^. Briefly, by treating the ligand as a translational degree of freedom that obeys the 1D Maxwell-Boltzmann distribution, the pre-factor can be approximated as (2πm/k_B_T)^-1/2^, where m is the mass of the di4using ligand.

## Acknowledgments

The VSC (Flemish Supercomputer Center) and the EuroHPC supercomputer LUMI provided the high-performance computing resources and services used for performing the simulations in this work. A.R.M. acknowledges research support from Ghent University (BOF starting grant no. BOF.STG.2021.0037.01).

